# Leveraging ancestry to improve causal variant identification in exome sequencing for monogenic disorders

**DOI:** 10.1101/010017

**Authors:** Robert Brown, Hane Lee, Ascia Eskin, Gleb Kichaev, Kirk E. Lohmueller, Bruno Reversade, Stanley F. Nelson, Bogdan Pasaniuc

## Abstract

Recent breakthroughs in exome sequencing technology have made possible the identification of many causal variants of monogenic disorders. Although extremely powerful when closely related individuals (e.g. child and parents) are simultaneously sequenced, exome sequencing of individual only cases is often unsuccessful due to the large number of variants that need to be followed-up for functional validation. Many approaches remove from consideration common variants above a given frequency threshold (e.g. 1%), and then prioritize the remaining variants according to their allele frequency, functional, structural and conservation properties. In this work, we present methods that leverage the genetic structure of different populations while accounting for the finite sample size of the reference panels to improve the variant filtering step. Using simulations and real exome data from individuals with monogenic disorders, we show that our methods significantly reduce the number of variants to be followed-up (e.g. a 36% reduction from an average 418 variants per exome when ancestry is ignored to 267 when ancestry is taken into account for case-only sequenced individuals). Most importantly our proposed approaches are well calibrated with respect to the probability of filtering out a true causal variant (i.e. false negative rate, FNR), whereas existing approaches are susceptible to high FNR when reference panel sizes are limited.

## Introduction

Vast decreases in the cost of exome sequencing have allowed for major advancements in the identification of causal variants for rare monogenic traits and disorders^1–4^. Although each individual carries 20,000-24,000 single nucleotide variants in their exome that differ from the human reference genome, most of these variants are common in the population or do not have a damaging effect and therefore are unlikely to explain a rare monogenic trait. Finding causal variants for monogenic traits through exome sequencing follows a two-step approach. First, variants that are too common to be consistent with the prevalence of a rare disorder are discarded^2^. Variants that remain under consideration are then prioritized based on frequency, functional, structural and conservation properties^5;6^, with more recent approaches using cross species comparisons^7^ or a combination of scores from several stand-alone methods or other data sources^8–11^. When pedigrees or cohorts of patients (with the same disorder) and their close relatives (e.g. parents or siblings) are sequenced, this two-step approach has proven to be extremely powerful in refining the list of prioritized variants to just a few variants^2–4; 12–16^. However, when only the case individual is available for sequencing, the number of variants that are left for follow-up in functional analysis is often on the order of hundreds of plausible variants ^10; 17; 18^, thus making it difficult to identify the causal variant(s).

In this work we present methods that leverage population structure (i.e. the variability in variant frequencies across populations) to improve the performance of exome sequencing studies of monogenic traits. Although it is commonly accepted in studying all types of disease that large well-matched control cohorts are important in limiting false positives^19^, monogenic disease studies often estimate variant frequencies across large databases of human variation at the level of continental ancestry (e.g. the Exome Variant Server^20^ European or African American data) instead of reference panels more finely tuned to the ancestry of the case individual(s); this is especially true when it is not practical to obtain a well-matched control group. Here, we investigate the use of matched allele frequency estimates (typically at the level of a country) to the ancestry of the sequenced individual^21; 22^. Since rare variants tend to be present in only a few closely related populations and absent from the rest^23–27^, the frequency estimates of alleles present in a given population will show a downward bias if estimated across individuals of multiple ancestries. That is, a variant might appear rare (<1%) across many populations, when in reality it is only rare in most populations and less rare or even common (>1%) in a few (see Supplemental Figure 1). As an example, consider variant rs17046386, it is generally rare or non-existent in non-Africans and present in Africans and those of African descent (see Figure 1). Based on European reference panels (or a global ancestry-unaware panel), this variant would not be excluded even though it’s relatively high frequency in Africans makes it unlikely to be pathogenic.

**Figure 1.**
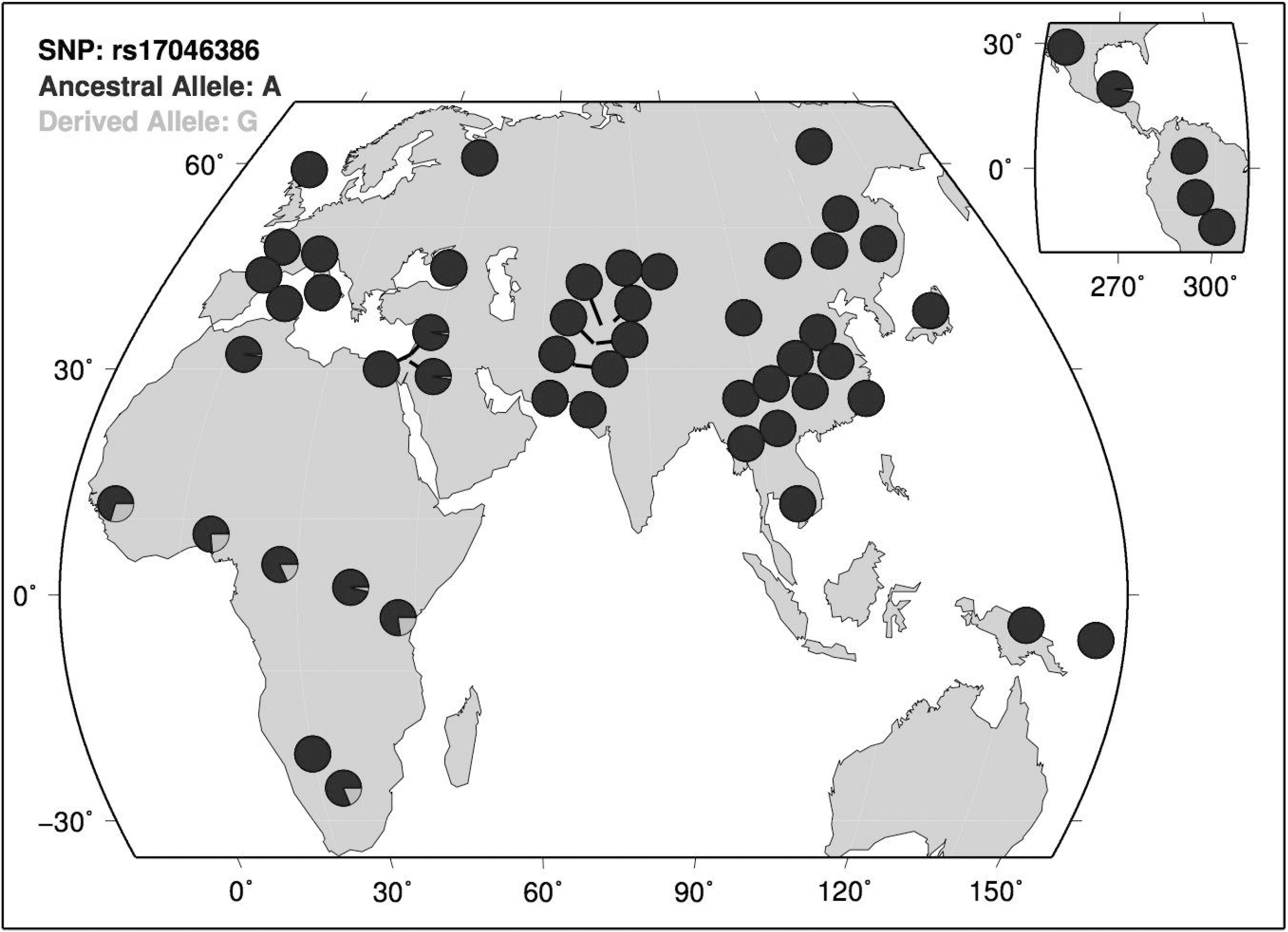
Geographic distribution of rs17046386 across the Human Genome Diversity Panel CEPH data. The minor allele is rare in non-African populations, but not rare in African populations.

The limited size of existing reference panels, especially when defining ancestry at the level of a country, induces significant statistical variance in the allele frequency estimates that must be accounted for. For example, a variant with a true frequency of 0.5% has an 8% chance of being observed with frequency >1% (and thus discarded) in a reference panel of 100 individuals as compared to only a 0.1% chance of being disregarded when frequencies are estimated over 1,000 individuals. To account for this effect, we introduce an false negative rate (FNR) filtering technique that protects against filtering a potential causal variant at a pre-specified significance level.

Starting from the 1000 Genomes^24^ and the NHLBI Exome Sequencing Project (ESP) Exome Variant Server (EVS)^20^ data sets, we use simulations to show that correctly matching reference panels to the country-level ancestry of the sequenced individual reduces the number of candidate causal variants from 418 to 351 in case-only simulations. Furthermore, by comparing frequencies across multiple populations, more variants can be removed (267 remaining if removing variants above the 5% FNR threshold in any population). In addition to case-only simulations, we simulate trio data for dominant and recessive diseases and show that ancestry aware filtering yields similar improvement under these scenarios. We validate our approach using exome-sequencing data from 20 real individuals with monogenic disorders for which the causal variants are known. In this data, without ever filtering out the true causal variant, our approach successfully reduces the mean number of heterozygous variants to be functionally tested from 750 to 604 (FNR<5%) when only matching to one population and to 435 (a 42% reduction) when leveraging all population data. Our results demonstrate that existing filtering pipelines for exome sequencing studies of monogenic traits can be significantly improved by taking ancestry into account. Finally, our results suggest that utilizing narrowly defined ancestry matched reference panels (i.e. at the country level) overcomes the reduction in performance due to higher statistical noise from the smaller panels.

## Methods

### Datasets

The 1000 Genomes Project^24^ has produced a public catalog of human genetic variation through sequencing individuals from different populations across the world. We use the 1000 Genomes individuals (1,092 in total) to evaluate the effectiveness of various filtering approaches (we removed the IBS from simulations due to the small number of individuals). Following the commonly accepted assumption that 85% of causal variants for monogenic traits are found in the exome^29^, we restrict our analysis only to variants found in the coding regions of autosomal chromosomes. For admixed individuals we matched ancestry locally using the local ancestry calls provided by the 1000 Genomes Project (the consensus calls of four different local ancestry inference methods^30–33^). Damaging scores for each single nucleotide variant were computed using the KGGSeq software with default parameters^10^ that combines the functional annotation scores in the dbNSFP^34^ database v2.0.

The NHLBI Exome Sequencing Project (ESP) Exome Variant Server (EVS) has released allele counts on 4,300 European-Americans and 2,203 African Americans^20^. PolyPhen2 scores are also provided for missense variants; the probably and possibly damaging predicted variants were used in the EVS-based analyses. We simulate different size reference panels using a binomial sampling with the allele frequencies estimated across all the European (or African American) EVS data as the true frequency.

To compare results seen in simulations to real data, we used exomes of 101 individuals with self-reported countries of origin including Turkey, Jordan, Tunisia, Egypt, Israel, Iran, Syria and Palestine. We grouped these individuals into a single supplemental reference population for estimating best matching allele frequencies. Of the 101 individuals, 20 had identified causal variants of monogenic disorders. Nine individuals were known to harbor heterozygous variants in genes causing autosomal dominant disorders, 10 individuals had homozygous variants, and one individual had potential compound heterozygous variants in genes causing an autosomal recessive disorder. We filtered out all variants except those with damaging annotations: splice acceptors, stop gains, frame shifts, stop losses, initiator codon changes, inframe insertions, inframe deletions, missense variants, splice region variants and KGGSeq predicted damaging variants. For consistency, the individual being examined was removed from the rest of the data that served as a reference panel. For comparison purposes, we included these individuals when estimating average frequencies across all populations in the 1000 Genomes project for the real data only.

### False negative rate estimation

We estimate the probability of filtering out a true causal variant (false negative rate) at a given frequency threshold as a function of a given reference panel and the maximum true allele frequency of the causal variant. The filtering threshold can be adjusted in order to provide a desired FNR. Let *t* be the nominal frequency threshold that is used for filtering. We define the corresponding FNR at this threshold as:

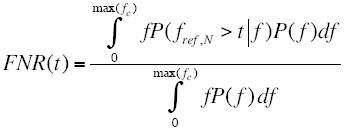

where *f* is the frequency of the variant in the population, max(*f*_*c*_) is the maximum assumed frequency of the causal variant in the population, P(*f*) is the proportion of variants with frequency *f* in the population and P(*f*_*ref,N*_ > *t*|*f*) is the probability that a variant with frequency *f* is observed at a frequency greater than *t* in the reference panel of *N* individuals randomly drawn from the population.

The FNR computation at a threshold *t* requires knowledge about the distribution of variants across all frequencies in the population; this can be estimated from population genetic theory under various demographic assumptions^25; 35–39^ or empirically from the data. In this work, we estimate the distribution P(*f*) from reference panel allele counts and perform the above integration across the observed site frequency spectrum as follows:

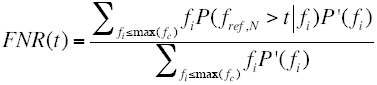

Here *fi* represents each of the unique allele frequencies observed in the reference panel of *N* individuals and P`(*fi*) represents the proportion of variants in the reference panel that have estimated frequency *fi*. The function sums over the discrete frequency values less than the max(*f*_*c*_). P(*fref,N* >*t* | *fi*) is modeled as a binomial draw with the frequency of success equal to *fi* and the number of draws equal to the number of allele counts (2*N*). Since the integration is over a discrete space we calculate the probability that the number of success is greater than the threshold times 2*N*. The max(*f*_*c*_) must also be larger than (2N)-^1^ for the FNR estimate to function. Having estimated a FNR for all possible filtering thresholds *t*, we propose to filter variants using the minimum frequency threshold *t* such that FNR(*t*)< 5%.

### Leveraging population structure for improved filtering

We compared different filtering approaches using simulated exomes from the 1000 Genomes data. For non-admixed individuals (e.g. not African-Americans) we use simulations where four haplotypes are drawn from a specific 1000 Genomes population and then paired to form two parental genomes. An offspring genome was then simulated from the parental genomes by passing variants according to Mendelian inheritance under the assumption that all sites are independent. We filtered out variants that do not result in an amino acid change or do not create or remove a stop codon. We compare three different scenarios likely to be encountered in a clinical diagnostic setting with 40 simulated offspring genotypes per scenario for each 1000 Genomes population (not including the IBS). The Case-Only scenario assumes there is no information on parental genotypes and there is no known mode of inheritance. The Trio-Dominant scenario assumes that both parental exomes are sequenced and the offspring and one of the parents has the disorder.

The Trio-Recessive scenario assumes that both parents are exome sequenced and heterozygous for the causal allele and that the offspring had two copies of the causal allele. We repeated the same analysis only using the variants for which there is a predicted damaging KGGSeq score (the variants in dbNSFP).

The cosmopolitan filtering approaches (*NoAncestry, f>1%* and *NoAncestry, FNR<5%)* estimate allele frequencies and FNRs across all 1000 Genomes individuals. The key intuition behind these approaches is that statistical noise is decreased with large reference panels at the cost of ignoring population structure. *NoAncestry, f>1%* filters out variants with allele frequency >1% without regard for the FNR, whereas *NoAncestry, FNR<5%* filters out variants above a threshold determined to ensure a desired FNR. We propose two ancestry-aware filtering approaches. The first (*PopMatched, FNR<5%*) uses only the reference individuals from the sub-continental population (country-level, see 1000 Genomes^24^) that best matches the sequenced individual. The second approach (*AllPop*) uses data across multiple sub-continental populations, by simply requiring that variants not be filtered out at each population’s FNR <5% threshold in each of a randomly chosen set of sub-continental populations. Sets sizes ranged between none and all the 1000 Genomes populations. The intuition behind this approach is that a variant common in at least one population is unlikely to be causal for monogenic disorders.

For admixed populations (MXL, PUR, CLM and ASW) we only assessed the Case-Only scenario and used the genotypes of the real admixed individuals from 1000 Genomes. In each individual at loci that are homozygous for African, European or Native American ancestry, we used continental allele frequency estimates obtained by averaging across the CEU, FIN, GBR and TSI for European frequencies, the YRI and LWK for African frequencies and the JPT, CHB and CHS for Native American frequencies. In local ancestry heterozygous regions we used a 50-50 weighting of the matching continental frequencies. In order to determine the FNR threshold, we first calculated the FNR threshold for the CEU, JPT and YRI and used the maximum threshold of those three populations. This is an overestimate of the true threshold because the allele frequencies will be downwardly biased and there is higher confidence in the allele frequency estimates due to larger reference panel sizes.

## Results

### Modeling statistical uncertainty increases filtering efficacy

We use simulations from the European-American Exome Variant Server (EVS)^20^ dataset to assess the increase in performance of FNR-based filtering as a function of panel size in a homogenous population. Figure 2a shows the threshold on the observed frequency as a function of reference panel size such that a FNR of 5% is maintained under different maximum frequencies of the causal variant (max(*f*_*c*_)). As expected, the frequency threshold that maintains a 5% FNR increases as reference panel sizes decrease (see Figure 2a). As the reference panel size is increased the filtering threshold that maintains a 5% FNR approaches max(*f*_*c*_).

**Figure 2.**
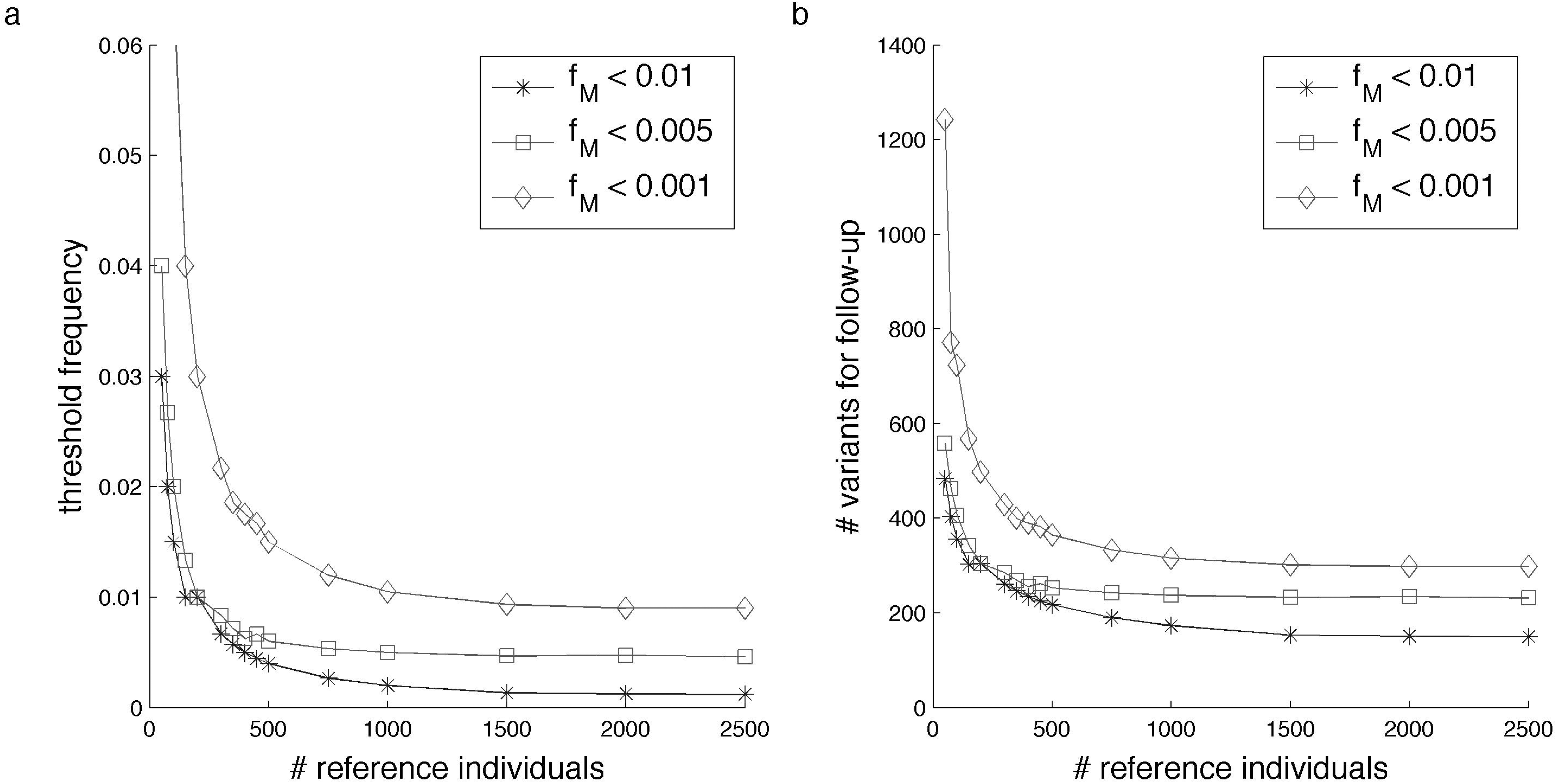
Reference panel size impacts the efficacy of filtering in exome sequencing in European simulations from the EVS data. Figure 2a shows the threshold on the variant frequency needed to achieve a 5% FNR for various assumptions about the maximum frequency of the causal variant in the population (from 0.001 to 0.01). Figure 2b displays the number of variants that remain to be followed up post-filtering at a 5% FNR rate. As expected with larger reference panel sizes, the estimated frequency from the reference panel becomes more accurate making the 5% FNR threshold converge to the maximum assumed frequency of the causal variant (fM) which in turn increases the efficacy of filtering. We observe limited gains in accuracy for reference panels over 500 individuals. Similar results are obtained for simulations of African Americans (see Supplemental Figure 2).

Figure 2b shows the average number of damaging or possibly damaging variants for follow-up (according to the EVS Polyphen2 ^40^ annotation scores) below the <5% FNR threshold. There is a diminishing return in filtering efficacy for reference panels larger than 500 individuals. This shows that as the assumed maximum frequency of the true causal variant (max(*f*_*c*_)) decreases, the 5% FNR threshold and number of variants for follow-up per individual also decrease.

Next, we investigated the effect of different maximum allele frequencies of the causal variant (max(*f*_*c*_)) and reference panel sizes on frequency-based and FNR-based filtering methods. Table 1 shows that for small reference panels (e.g. ∼100 reference individuals, approximately the size of a 1000 Genomes country-level population) the frequency-based approach is mis-calibrated with respect to the probability of filtering out the true causal variant. Although the approach that maintains a proper FNR<5% significantly increases the number of variants for follow-up from 298.0 to 724.1 on average, this is necessary as it reduces the FNR from 25% for the frequency-based approach to 5% for the FNR-based approach. In contrast, when reference panels are large, the frequency-based approach is too conservative (due to the lack of a FNR calculation) leading to an increased number of follow-up variants. For example, the FNR<5% approach reduces the number of variants for follow-up from 311 to 150 variants, under the assumption that the max(*f*_*c*_) is 0.1% and 2500 reference individuals. Qualitatively similar results were observed when simulating exomes from the EVS African-American data (see Supplemental Figure 2).

**Table 1.** Method comparisons for different reference panel sizes and maximum causal allele frequencies. We compare two methods. The first is a method (f > 1%) that filters out any variants at an observed frequency >1% ignoring the statistical noise on the frequency estimates (and thus the FNR). The second is a method (FNR<5%) that filters out variants if observed above a threshold frequency guaranteed to provide less than a 5% chance of filtering out the true causal variant. At small reference panel sizes it is critical to incorporate statistical noise from the reference panel to not over-filter the true causal variants. Conversely, with large reference panels, a hard 1% frequency filter is too conservative and significantly increases the number of variants remaining for follow-up analysis.

|  |  | 100 Reference Individuals |  |  | 2500 Reference Individuals |  |  |
| --- | --- | --- | --- | --- | --- | --- | --- |
| Max True Frequency | Method | Threshold | Number of Variants for Follow-up | Probability of Filtering True Causal | Threshold | Number of Variants for Follow-up | Probability of Filtering True Causal |
| 1.00% | (f > 1%) | 1.0% | 298.0 | 25.4% | 1.0% | 310.9 | 2.2% |
|  | FNR<5% | 6.5% | 724.1 | 4.6% | 0.9% | 298.3 | 4.8% |
| 0.10% | (f > 1%) | 1.0% | 298.0 | 12.1% | 1.0% | 310.9 | 0.0% |
|  | FNR<5% | 1.5% | 356.0 | 4.3% | 0.1% | 149.8 | 3.6% |
| 0.05% | (f > 1%) | 1.0% | 298.0 | 8.0% | 1.0% | 310.9 | 0.0% |
|  | FNR<5% | 1.5% | 356.0 | 1.9% | 0.1% | 141.2 | 2.3% |

### Leveraging ancestry to increase filtering performance

We next assessed the performance of filtering with or without including ancestry information to account for the highly structured nature of rare variants^23–25; 41^ in simulations of non-admixed individuals starting from the 1000 Genomes data. We compared several methods for filtering variants in exome studies under Case-Only, Trio-Dominant and Trio-Recessive scenarios. Under all disease architecture and trio scenarios the methods that take ancestry into account outperform methods that do not (Table 2). The *PopMatched, FNR <5%* and *NoAncestry, FNR <5%* each filter based on a threshold determined to ensure a FNR of at most 5%. The difference is that the *NoAncestry* method uses the entire 1000 Genomes data set as a single reference population whereas *PopMatched* uses only the 1000 Genomes individuals from the same population as the simulated case individual as references. The population matching decreases the number of variants by at least 15% under the Case-Only, Trio-Dominant and Trio-Recessive scenarios; this is true when only using KGGSeq predicted damaging variants as well (Table 2). This improvement comes regardless of the fact that the <5% FNR filtering threshold in the *PopMatched* method (average filtering threshold is 2.2% and always >2%) is on average twice that of that *NoAncestry* method (filtering threshold is 1.1%) due to the significantly reduced reference panel size (average of 93 individuals per 1000 Genomes population). This demonstrates that the benefit of better population matching outweighs the cost of higher statistical noise resulting from the small reference panels. The Trio-Dominant scenario has approximately half as many variants for follow-up as the Case-Only scenario (288 compared to 582 for the *PopMatched, FNR<5%* method, Table 2), this would be expected assuming one parent were also affected. The Trio-Recessive scenario, simulated without inbreeding, shows very few variants for follow-up (<6 under all scenarios and filtering methods), this is expected when looking for rare variants appearing homozygotically in an individual (Table 2).

**Table 2.** Average number of variants that remain for follow-up post-filtering in simulations of non-admixed individuals. All FNR approaches assume the maximal causal variant frequency of 1%. Poorly matched reference panels greatly affect the number of variants for follow-up analysis more so than accounting for increased statistical error from smaller reference panels. The top method, 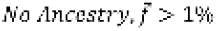 has an increased FNR (6.0%) relative to all the other methods. Case-Only represents filtering with exome data from a single individual and makes no assumptions about disease architecture. Trio-Dom assumes the case individual and both parental exomes are sequenced and that only one parent has the dominant disorder. Trio-Rec assumes there is exome data from the case individual and both parents and that both parents are carriers of one copy of the causal variant but do not have the recessive disorder.

|  | Number of variants for follow-up |  |  | Number of variants for follow-up, with KGGSeq variants |  |  |
| --- | --- | --- | --- | --- | --- | --- |
|  | Case-Only | Trio-Dom | Trio-Rec | Case-Only | Trio-Dom | Trio-Rec |
| *NoAncestry, $f > 1\%$* | 678.9 | 334.7 | 5.3 | 409.5 | 202.4 | 2.9 |
| *NoAncestry, FNR < 5%* | 695.7 | 342.7 | 5.6 | 418.2 | 206.5 | 3.1 |
| *PopMatched, FNR < 5%* | 581.9 | 288.2 | 3.1 | 350.6 | 173.8 | 1.7 |
| *AllPop* | 415.9 | 207.8 | 1.6 | 267.3 | 133.3 | ~~5.40~~ |

We observe that the *NoAncestry, FNR <5%* approach leads to a slightly increased number of variants that need to be functionally followed up over the *NoAncestry, f>1%* approach (Table 2). The increase in the number of variants is necessary to attain a correct 5% FNR rate (*NoAncestry, f>1%* attains a FNR of 6.0%).

Finally, we assessed a method that filters variants observed in any population above each population’s 5% FNR threshold in addition to the best-matched population (*AllPop*). All FNR-based approaches assume that max(*f*_*c*_) <1% in all populations. The *AllPop* approach does not account for multiple testing, but does demonstrate that as more population are randomly sampled, many variants can be eliminated due to high frequency in some populations that are inconsistent with causal allele frequency assumptions. This method shows a 40% reduction from the *NoAncestry, FNR<5%* method in the Case-Only scenario (Table 2). In Figure 3, we investigated how the number of follow-up variants decreases as a function of the number of reference populations available in the Case-Only scenario. With data from more populations available, there is a greater chance of observing that a given variant is common in at least one population, and is thus unlikely to cause a rare disorder. This demonstrates that there are a significant number of variants that are common in some populations but rare in the population of the case individual. Careful selection of a few populations genetically distant from a case individual’s population can further reduce the number of variants remaining beyond what is obtainable through only the *PopMatched, FNR<5%* method and will reduce the effects of multiple testing.

**Figure 3.**
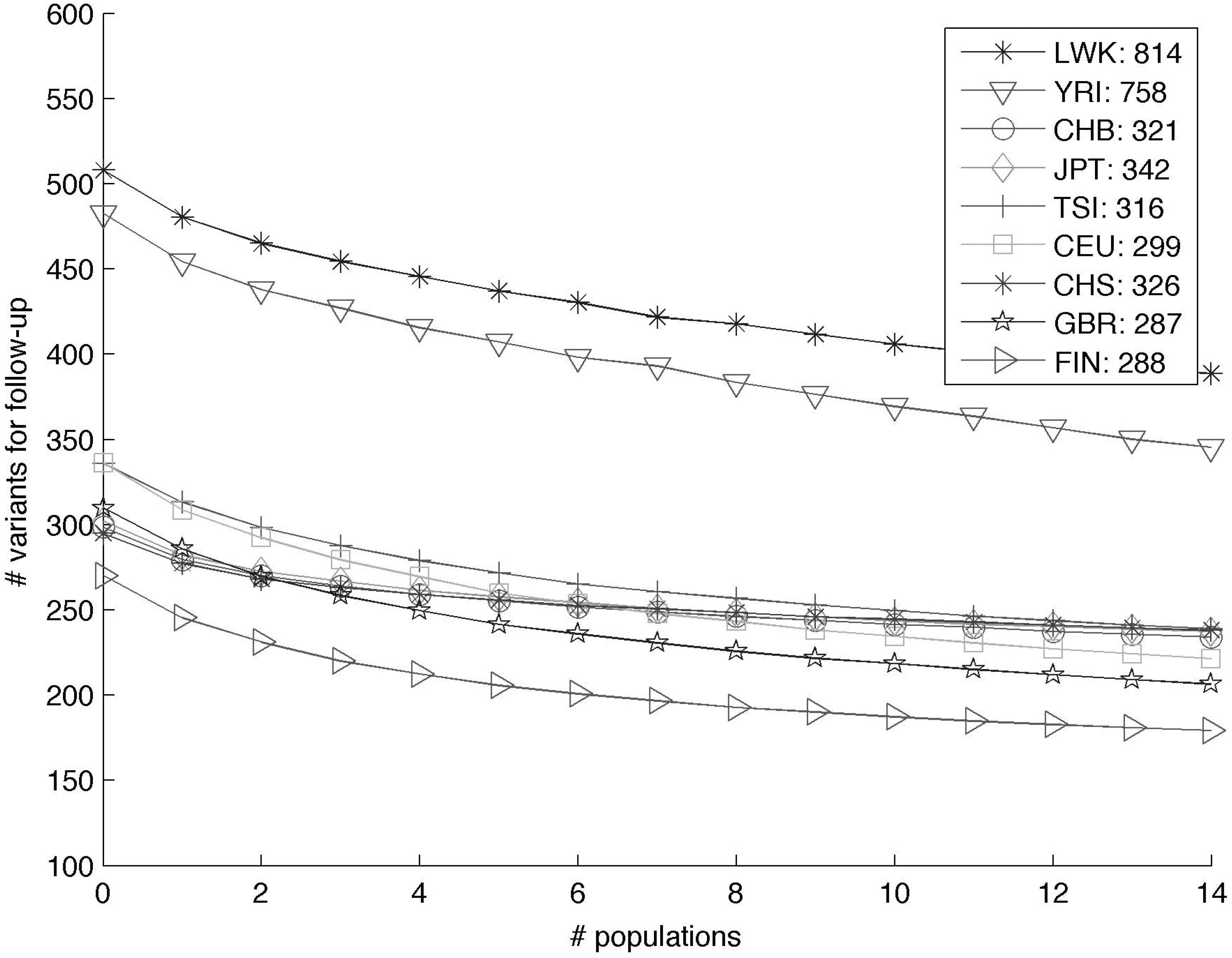
Estimates of the number of case-only variants for follow-up for the *AllPop* method for various numbers of additional comparison populations. Starting form the results of the *PopMatched, FNR < 5%* approach (shown at x-axis=0 in the plot) additional randomly chosen populations are added to the reference panels. Increasing the amount of information about various populations can further reduce the number of variants for follow-up analysis thus increasing the efficacy of filtering.

Finally, we observe a similar pattern of improved performance when restricting to functionally damaging variants as predicted by KGGSeq (Table 2). Taken together, this shows that in all the aforementioned scenarios the improvements of ancestry-aware filtering do not come exclusively from variants that would have been filtered later due to non-damaging predictions, but rather, from variants removed proportionally from damaging and non-damaging predicted variants.

### Ancestry-aware filtering in admixed individuals

Results above were obtained using individuals of homogeneous ancestry. We extend our approach to populations of admixed ancestry (e.g. African Americans) by considering their local ancestry structure. Individuals with recent ancestry from multiple continents have genomes that are a mosaic of segments each originating from different ancestral populations. We incorporate the local ancestry structure in the filtering step with the *PopMatched-LA, FNR <5%* approach that matches reference panels by ancestry according to each site in an individual’s genome (see Methods). This significantly lowers the number of variants for follow-up in the admixed populations as compared to the local ancestry naïve method (*PopMatched, FNR <5%)* (see Figure 4). When using information from all populations in the 1000 Genomes dataset, there is improvement for all admixed populations over the method that ignores ancestry (*NoAncestry, FNR <5%)* (see Figures 4 and Supplemental Figure 3). For example, in African American individuals we observe a reduction from 668 to 592 variants from just matching the local ancestry to continental populations as compared to using all 1000 Genomes data with a FNR <5%. There is significant diversity within subpopulations of Mexico that may make finding well-matching reference panels difficult^42^. While correct matching has been shown to be very beneficial, incorrect matching may increase the number of variants remaining for follow-up. This could explain why the *PopMatched-LA, FNR <5%* approach performs worse for the MXL than the *PopMatched, FNR <5%* approach.

**Figure 4.**
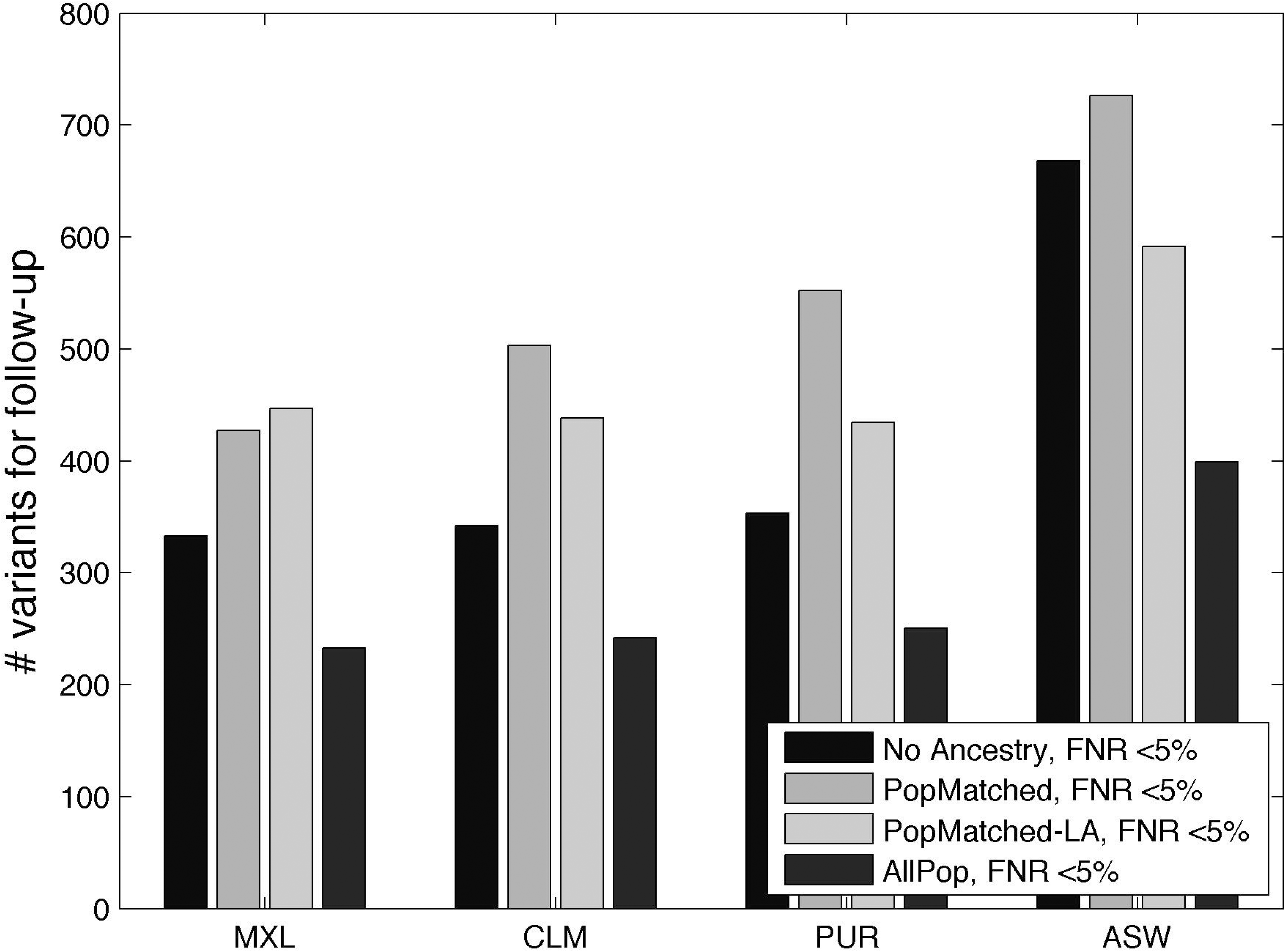
Population matching using local ancestry information improves performance over local ancestry naïve population matching in admixed populations. The MXL has worse local ancestry aware performance as compared to local ancestry naïve performance likely due to reference panels being poorly matched given the high levels of diversity in Mexicans.

### Analysis of 20 exomes of individuals with monogenic traits

To examine the performance of the different filtering strategies when applied to actual data, we used the data from 20 real exome sequenced individuals with monogenic disorders where the causal variants have been previously identified. We assumed a maximum causal allele frequency of 1% for all cases because there was no prevalence data^2^. For all modes of inheritance, the number of variants in an individual for follow-up after filtering was lower when filtering with the *PopMatched, FNR <5%* and *AllPop* approaches as opposed to the *NoAncestry, f>1%* approach that does not account for the FNR (See Supplemental Table 1). For example, using our approach only 604.8 (434.6 if using all populations’ data) variants need to be followed-up for a dominant disorders as compared to 749.7 for the *No Ancestry, f>1%* approach (see Table 4). Importantly, for all of the filtering schemes, the true causal variant identified in these real individuals was never filtered out. This demonstrates population matching for filtering allele frequencies significantly reduces the number of variants remaining for follow-up analysis, while still maintaining an appropriate false negative rate. It also shows that when researchers can safely assume a causal allele frequency <1% in many populations, they can substantially further reduce the number of variants remaining for follow-up analysis. This real data analysis shows the importance of being able to first filter with a well-matched reference panel, even of just 100 individuals. In Supplemental Table 1 we report the variants remaining in each individual along with country data, reported inheritance pattern, the zygosity of the causal variant and assumed disorder architecture.

## Discussion

Exome sequencing for rare monogenic disorders has proven to be very useful for discerning the causal genes for these traits. Although extremely powerful when closely related individuals are simultaneously sequenced, current pipelines for exome sequencing of a single individual often yield too many variants to be tractable for functional follow-up. In this work, we introduce approaches that account for the finite sample size of the existing reference panels used in filtering while jointly modeling the ancestry to improve the filtering step. Both the real data analysis of 20 exomes of individuals with known monogenic disorders and the simulations show that our approaches reduce the number of variants that need to be further investigated, thus increasing the effectiveness of identifying causal variants using exome sequencing of unrelated individuals. This work demonstrates that in a clinical setting, even a small reference panel of 100 individuals from a well-matched population can have significant impact on the filtering ability of a case individual.

The current bottleneck in using population structure to help identify rare variants is the limited size of the reference panels for specific narrowly defined populations. Current reference panels show that at the 1% allele frequency level that there are significant differences between populations. However, it is difficult to assess allele frequencies below this level due to reference panel size for specific populations. This demonstrates the need for larger reference panels of more narrowly defined populations (not just continental level) in order to fully assess the structure of rare variation. As the panel sizes increase, researchers studying monogenic disorders will be able to use smaller maximum causal allele frequencies when appropriate for the trait of interested. Furthermore, the larger reference panels will also lower filtering thresholds.

With the increasing availability of public databases it may be possible to obtain accurate estimates of disorder prevalence across populations. Our proposed approaches could be further extended to take this information into account by using different thresholds on the maximum frequency of causal variants (max(*f*_*c*_)) across populations. We leave that as ongoing and future work.

**Table 3.** Different levels of genetic diversity across populations induce a variation in the average number of variants remaining for follow-up in an individual. The highest number of variants remaining for follow-up is seen in African populations (YRI and LWK) as well as African-Americans (ASW); this is consistent with these populations have the greatest amount of genetic diversity. * denotes admixed populations where a local ancestry aware method was utilized (see Methods).

| 1000 Genomes<br>Population<br>(Number of<br>Individuals) | *NoAncestry*,<br>*FNR* <5%, (s.d.) | *PopMatched*,<br>*FNR* <5%, (s.d.) | *AllPop*, (s.d) |
| --- | --- | --- | --- |
| ASW* (61) | 668.2 (75.9) | 591.5 (39.9) | 398.8 (40.2) |
| CEU (85) | 298.9 (32.4) | 336.2 (36.2) | 221.4 (26.7) |
| CHB (97) | 320.7 (31.4) | 298.7 (34.2) | 234.0 (27.4) |
| CHS (100) | 326.8 (18.2) | 295.0 (24.1) | 238.1 (15.2) |
| CLM* (60) | 342.0 (39.2) | 438.0 (40.0) | 242.0 (20.5) |
| FIN (93) | 288.5 (25.1) | 270.2 (36.9) | 179.3 (21.2) |
| GBR (89) | 286.6 (27.0) | 309.8 (33.1) | 206.4 (22.5) |
| JPT (89) | 342.1 (22.6) | 302.0 (25.9) | 237.4 (18.5) |
| LWK (97) | 814.5 (38.3) | 507.9 (36.6) | 388.7 (29.0) |
| MXL* (66) | 332.9 (25.1) | 446.6 (37.2) | 232.7 (22.6) |
| PUR* (55) | 353.3 (48.5) | 434.3 (44.9) | 250.2 (29.0) |
| TSI (98) | 315.6 (24.6) | 336.0 (26.7) | 238.9 (21.4) |
| YRI (88) | 757.7 (29.9) | 482.5 (33.5) | 345.4 (20.6) |

**Table 4.**
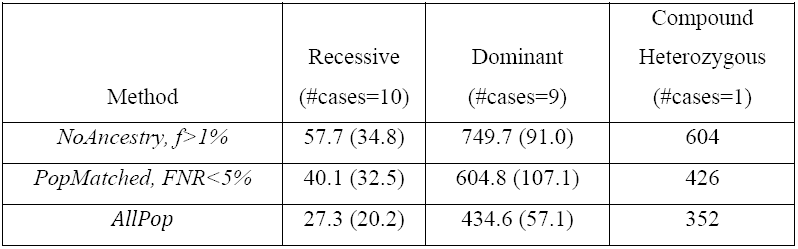
Average number of variants that remain for follow-up post-filtering in real exome studies of 20 individuals with Mendelian disorders. None of the filtering approaches removed the true casual variants from consideration. Across all disorder architectures, we observe a significant decrease in the number of variants that need to be followed up if ancestry is incorporated in the filtering step. Parentheses denote standard deviations. Variants were eliminated from consideration as potentially true causal variants if they are not annotated as damaging (see Methods) and if they are not observed twice if the disorder is assumed to be autosomal recessive or at least once if it is assumed to be dominant (heterozygous) or compound heterozygous.

## Supplemental Data

The supplemental data contains three figures and one table.

## Acknowledgements

This work is supported in part by the National Institutes of Health (R03-CA162200, R01-GM053275 to BP and T32-HG002536 to RB). AE and SFN are supported by the Genomics/Informatics Core of the UCLA MUSCULAR DYSTROPHY CORE CENTER from NIAMS (P30AR057230). B.R. is a fellow of the Branco Weiss Foundation and an A*STAR and EMBO Young Investigator. This work was funded by a Strategic Positioning Fund for Genetic Orphan Diseases and an inaugural A*STAR Investigatorship from the Agency for Science, Technology and Research in Singapore. The funders had no role in study design, data collection and analysis, decision to publish, or preparation of the manuscript.

### Web Resources

We provide publicly available software implementing our approach: http://bogdan.bioinformatics.ucla.edu/software/

Figure 1 was generated using the following website with data from the Human Genome Diversity Panel^43; 44^: http://hgdp.uchicago.edu/cgi-bin/gbrowse/HGDP/

